# Pollen diet, more than geographic distance, shapes provision microbiome composition in two species of cavity-nesting bees

**DOI:** 10.1101/2025.03.17.643742

**Authors:** Rachel L. Vannette, Neal M. Williams, Stephen S. Peterson, Alexia N. Martin

**Affiliations:** Department of Entomology and Nematology, University of California Davis, Davis, CA, USA; Foothill Bee Ranch, Foresthill, CA, USA

**Keywords:** food microbiome, intraspecific variation, host-microbe interactions, *Osmia*, solitary bee

## Abstract

The microbial composition of stored food can influence its stability and determine the microbial species consumed by the organism feeding on it. Many bee species store nectar and pollen in provisions constructed to feed developing offspring. Previous work has shown variation in provision microbiome among bee populations, yet whether this variation is determined by the pollen types within provisions, variation between bee species at the same nesting sites, or geographic distance was unclear. Here, we sampled two species of co-occurring cavity nesting bees in the genus *Osmia* at 13 sites across the Sierra foothills in California and examined the composition of pollen, fungi and bacteria found in their provisions across sites. As expected, pollen, bacterial and fungal composition exhibited significant turnover between bees and sites, with bee species characterized by particular pollen and microbial species. Pollen composition explained 15% of variation in bacterial composition and ∼30% of variation in fungal composition, whereas spatial distance among sites explained minimal additional variation. Symbiotic or bee-specialized microbe genera *Ascosphaera*, *Sodalis* and *Wolbachia* showed contrasting patterns of association with pollen composition, suggesting distinct acquisition and transmission routes for each. Comparing provisions from both bee species comprised of the same pollens points to environmental acquisition rather than bee species as a key factor shaping the early stages of the bee microbiome in *Osmia*. The patterns we observed also contrast with *Apilactobacillus*-dominated provision microbiome in other solitary bee species, suggesting variable mechanisms of microbial assembly in stored food among bee species.

## Introduction

Many animal species store food for later consumption or as provisions for developing offspring. The microbial populations within stored foods can be a key factor in food stability (Louw *et al*. 2023; Paul Ross *et al*. 2002) and can contribute to the gut microbiome of consuming organisms (Stothart *et al*. 2023). The microbiome of stored foods of a species often varies with geography. Such variation is documented in the seed caches of woodrats (Herrera *et al*. 1997) and fungal gardens of leaf cutter ants, as well as in human foods like spontaneously fermented wine (Li *et al*. 2022; Morrison-Whittle & Goddard 2018) and lactic acid-driven vegetable fermentation (Miller *et al*. 2019). However, many factors vary geographically, including components of provisions (e.g., plant species used by woodrats or ants), storage and processing conditions, and environmental microbial communities from which food microbes originated (e.g., soil, water or plant surfaces). As a result, the contribution of each factor in driving variation in microbiome composition is often difficult to disentangle.

Adult females of many bee species store nectar and pollen that is consumed by developing offspring. Solitary bees–species with no cooperative brood care, division of reproductive labor or communal foraging–construct brood cells in wood, pithy stems, soil or other cavities. Brood cells are typically provisioned with enough nectar and pollen to support the development of single offspring (Danforth *et al*. 2019). After completing the provision, a female lays an egg on it and seals the brood cell. Most species have no interaction with food or their offspring after the brood cell is sealed (offering no opportunity for continued maintenance of stored food). Eggs typically hatch within 3-10 days and the growing larvae consume the stored provision over a few weeks. Thus, the carbohydrate, lipid and protein-rich provision remains in cavities for days to weeks, allowing ample time for microbial growth and community dynamics. Despite adaptations of bees to reduce microbial growth within these stored larval food provisions (Burgett 1974, Wilson, et al 2015), provisions are commonly colonized by bacteria and fungi (Gilliam *et al*. 1989; McFrederick, Quinn S. *et al*. 2012; Steffan *et al*. 2024) largely sourced from the environment (Anderson *et al*. 2013).

For solitary bees, emerging evidence suggests that the microbiome of stored food can affect multiple aspects of bee fitness. Mortality resulting from fungal infection of developing bees and their stored food (Aronstein 2013) is thought to be a major driver of bee population dynamics, particularly for non-social bees (Minckley & Danforth 2019). The incidence of failed nests due to fungi or bacteria in solitary bees ranges from 20-70% depending on bee species and environmental conditions (Cane *et al*. 1983). Antagonistic microbes include bee-specialized species like some *Ascosphaera* (L.S. Olive & Spiltoir, 1955) (Onygenales: Ascosphaeraceae; Wynns 2012) that grow uniquely on the stored provisions of bees or opportunistic pathogens including *Aspergillus* (Evison & Jensen 2018). However, bacteria and fungi are common in healthy brood cells and these nonpathogenic microbes have been hypothesized to preserve or transform stored nectar and pollen (Anderson *et al*. 2014; Gilliam *et al*. 1988; Hammer *et al*. 2023; Steffan *et al*. 2024). In some cases microbes can supplement bee nutrition or synthesize growth-limiting resources (Paludo *et al*. 2018, Dharampal *et al*. 2019). Microbial communities may also suppress the growth of bee pathogens (Christensen *et al*. 2023; Voulgari-Kokota *et al*. 2020) via antimicrobial activity (Cambronero-Heinrichs *et al*. 2019). In solitary bee provisions sampled to date, the microbiome composition has been shown to vary among bee species (Voulgari-Kokota *et al*. 2019), pollen source and geographic locations (Dew *et al*. 2020; McFrederick & Rehan 2019; Nguyen & Rehan 2023; Rothman *et al*. 2020), yet the degree to which bee species traits, differences in pollen types used, or other environmental variables drive such patterns is unclear.

Here, we examine fungal and bacterial communities in brood cell provisions among populations of two species of solitary bees within the genus *Osmia.* We compare whether pollen composition or geographic proximity among populations better predicts microbial composition in provisions, both within and across bee species. We also compare factors predicting variation in the presence and abundance of commonly insect-associated symbiotic bacteria (e.g. *Sodalis*) and or bee-associated fungi (e.g. *Ascosphaera*) which may differ from other microbes in their acquisition and transmission routes. We hypothesize that if local geographic factors drive microbial composition in stored food, microbial community composition should be increasingly different with geographic distance regardless of bee species. On the other hand, if bee species identity most strongly determines microbial community of bee provisions, we predict microbial communities will differ between two species sampled at the same site and within a species, microbial communities may vary less among geographic sites. Finally, if both pollen type and bee species determine microbial composition, then we expect strong differences between the two bee species, which differ in dominant pollen used but also differences within each species associated with shifts in pollen collection among brood cells.

## Methods

### Study system

We sampled the stored nectar and pollen within brood cells of two species of cavity-nesting *Osmia* (mason bees, Megachilidae) that nest in early spring with partially overlapping phenology, and that often co-occur. It is common to see nests of the two species within the same piece of wood nesting substrate within a microsite. *Osmia lignaria* is managed as a pollinator for early-flowering orchard crops and is a diet generalist (Williams & Tepedino 2003). *Osmia ribifloris* is a diet specialist that forages mainly on host plants in the Ericaceae (Sampson *et al*.

2013). Both *Osmia* species are common across the western United States. Nests of each species are distinguished by nesting material: *O. lignaria* constructs nest partitions using mud while *O. ribifloris* uses masticated leaves (Fig 1).

**Figure 1.**
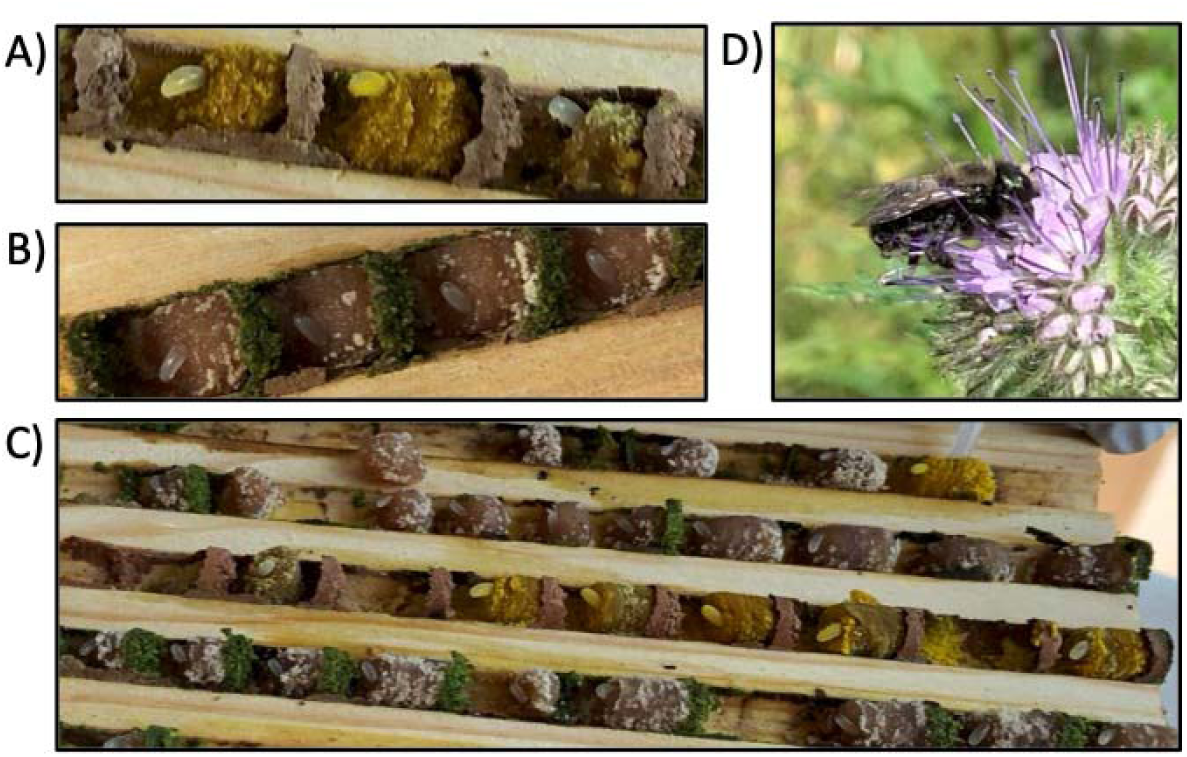
*Osmia* bee nest structure. *Osmia lignaria* brood cells (A) and *O. ribifloris* brood cells (B) are shown in grooved wooden boards. Linear nests of each species (C) typically contain between 6-10 brood cells separated by partitions, each with an individual egg. *Osmia lignaria* constructs partitions using mud while *O. ribifloris* uses masticated leaf material. At each site and for each species, a maximum of three nests were sampled, with two brood cells sampled per nest. (D) Adult *Osmia lignaria* foraging on *Phacelia* flower. Photos by N. Williams and R. Vannette.

### Sampling design

We sampled *Osmia* populations at 13 geographic locations, hereafter ‘sites’, in the foothills of the western slope of the Sierra Nevada in California (Fig 2) where populations of each species were established or bees were newly released. At each site, nesting was encouraged by the placement of clean (washed and heat-treated) nesting blocks in early spring. In 2020, populations were sampled at 10 sites, where sites hosted natural populations of either or both species (Table S1). In 2022, populations of *O. ribifloris* were released at 3 agricultural sites and a subset of the natural source populations were also re-sampled (Table S1).

**Figure 2.**
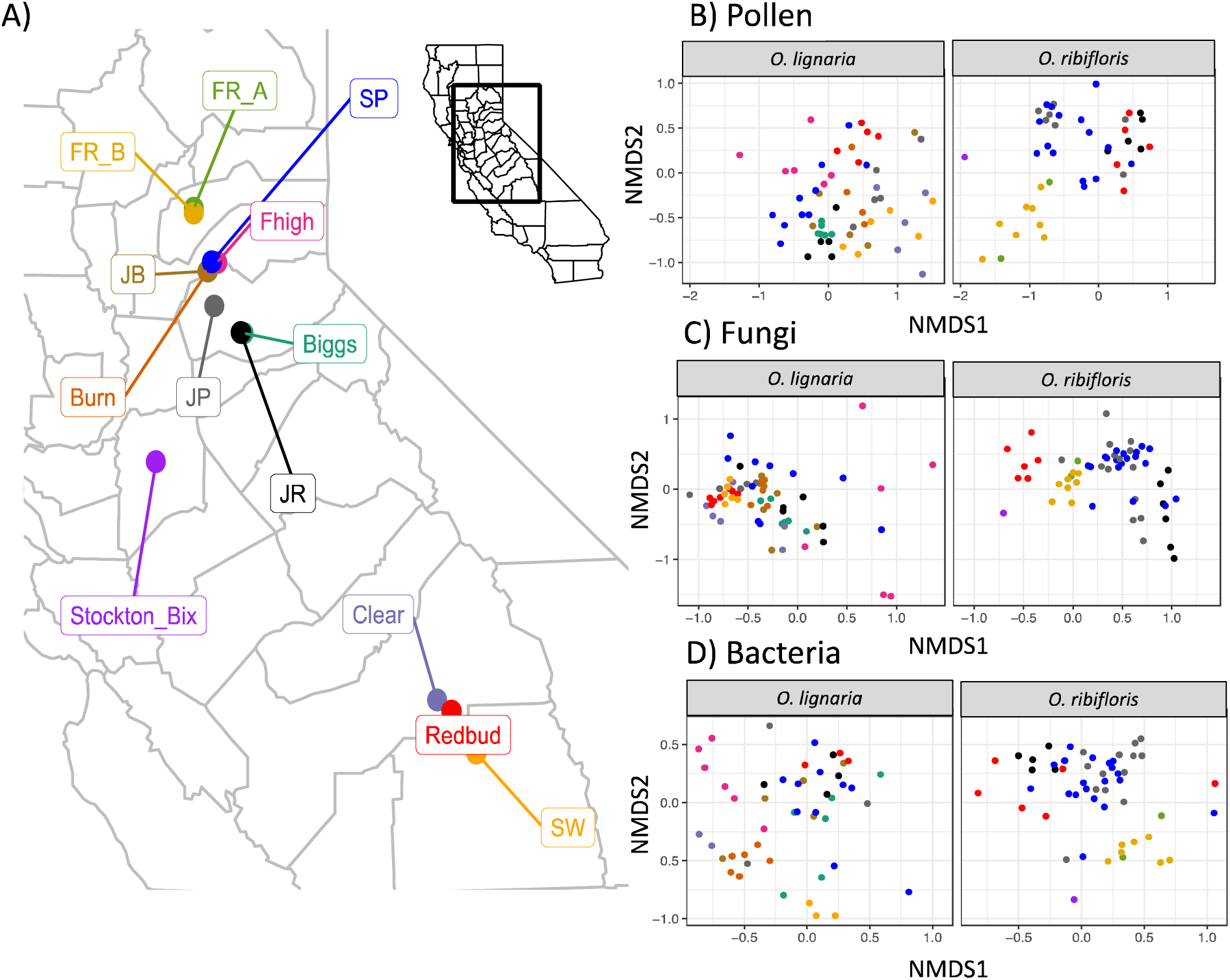
Map of bee sampling locations (A) and variation in provision composition among sites using NMDS (Non-metric Multidimensional Scaling) Plots of A) pollen composition (based on Jaccard dissimilarity), B) fungal (ITS) community composition and C) bacterial (16S) composition based on amplicon sequencing, visualized among bee species *Osmia lignaria* (L) and *Osmia ribifloris* (R) and sites. Ordinations were performed on a dataset including both species together but are faceted by species for clarity. Provision samples are colored by sampling site. Both species could be sampled at 4 sites, while *O. lignaria* were sampled at an additional 6 sites and *O. ribifloris* were present at an additional 3 sites (Table S1).

To collect provision material, we transferred whole provisions to sterile 1.7 mL centrifuge tubes using single use plastic pipets or sterile toothpicks. Provisions were sampled when brood cells contained either an egg (most common), 1^st^ or rarely 2^nd^ instar larvae, and no frass was present in brood cells. Larvae of these species have a blind gut and do not defecate until the 5^th^ instar (Torchio 1989). Collected provisions were stored at -80°C until DNA extraction. At each site, we sampled provisions from 3 nests per species and 2 brood cells within each nest, or as many as were available.

### Sample Processing

DNA was extracted from approximately a third of each provision using the Qiagen PowerSoil kit (Germantown, MD, USA) following manufacturer’s instructions. Extracted DNA was quantified using nanodrop and submitted for amplicon sequencing at Dalhousie Integrated Microbiome Resource (IMR, Dalhousie University, Nova Scotia CA) Sequencing facility. Plant sequences were targeted using rbcL primers rbcLaF (5’-ATGTCACCACAAACAGAGACTAAAGC-3’) and rbcL506R (5’-AGGGGACGACCATACTTGTTCA-3’), bacterial 16S rRNA sequences were amplified using 799F (799F=5’-AACMGGATTAGATACCCKG-3) and 1115R (1115R=5’-AGGGTTGCGCTCGTTG-3’) to reduce contamination with mitochondrial and chloroplast sequences and fungal ITS sequences were amplified using ITS86 (5’-GTGAATCATCGAATCTTTGAA) and ITS4 (5’-TCCTCCGCTTATTGATATGC-3’). Kit extraction blanks were also included in sequencing runs for all sample types (N=2/region). PCR was performed in duplicate using high-fidelity Phusion Plus polymerase, followed by amplicon cleanup and normalization using Charm Just-a-Plate Normalization kit, and sequencing using MiSeq PE300 was performed by Dalhousie IMR.

### Sequence data processing

Sequence data were quality filtered, trimmed, error-corrected, merged, and had chimeras removed using the DADA2 pipeline (Callahan *et al*. 2016), see full code in accompanying Dryad and Zenodo repository. This generated a table of actual sequence variants (ASVs) present across each sample. Sequence variants detected in the negative controls were removed from the dataset using Decontam (Davis *et al*. 2018), resulting in the removal of 8 fungal ASVs, 3 pollen ASVs and 14 bacterial ASVs. Taxonomic annotation was performed using the UNITE database (v7.2) for ITS sequences (Nilsson *et al*. 2019), BEExact v2023.01.30 and SILVA v138.1 for 16S sequences, and plant database for rbcL sequences (Bell & Brosi 2016). Manual BLAST searches were performed for the most abundant 100 ASVs in each dataset when unannotated at the family level, or when 16S databases did not match, and updated when BLAST searches indicated high confidence annotations. Data wrangling was performed using Phyloseq (McMurdie & Holmes 2013). All ASVs that were not assigned to an appropriate kingdom based on the sequencing region (e.g. fungi for ITS) and a phylum within that kingdom were filtered out; reads assigned as mitochondria or chloroplast were also filtered out from ITS and 16S datasets. Following sequence curation, species accumulation curves were examined to ensure saturation (Figure S1), with samples dropped below a minimum of 500 reads for pollen (mean = 25,110, 2 samples dropped), 450 reads/sample for fungi (mean=11,564, none dropped), and 500 for bacteria (mean=6,570; 20 samples dropped).

### Statistical analyses

Statistical analyses were performed using R v.4.3.1 (R Core Team 2021). We compared the taxonomic alpha diversity and community composition of pollen (rbcL), bacteria (16S) and fungi (ITS) between bee species and among populations (geographic site). For fungi and bacteria, reads were normalized within a sample for community and diversity analyses (McMurdie & Holmes 2013). First, we examined if alpha diversity of pollens, bacteria or fungi varied among species or sites. We estimated alpha diversity of pollen in provisions using the Chao1 estimator implemented in Phyloseq at the genus level and at the ASV level for bacteria and fungi. We examined variation in species richness for each group using linear models with bee species, site and their interaction as predictors. Then, we tested whether genus-level diversity (Chao1) of pollens in a provision predicted the diversity (Chao1) of bacterial or fungi using linear mixed models, with nest ID as a random effect.

We also examined predictors of beta diversity in pollen, bacterial and fungal composition. Bray-Curtis dissimilarity was used for bacterial and fungal community analyses. For pollen communities we used Jaccard dissimilarity indices, which do not consider differences in relative abundance, because pollen metabarcoding sequence abundance can deviate substantially from proportional pollen relative abundance (Bell & Brosi 2016). Community dissimilarity was visualized using PCoA and the degree to which species, site ID or their interaction predicted dissimilarity in community composition was assessed using perMANOVA implemented using adonis2 in vegan (Oksanen *et al*. 2007).

To examine which bacteria, fungi or pollens at the genus level distinguish bee species or sites within a bee species, we conducted an analysis of differential abundance using DESeq2 (Love *et al*. 2014). To test if bacterial or fungal composition of the provision can be used to predict bee species that constructed it, we used a random forest approach (Liaw & Wiener 2002) to estimate classification error rate and rank genera by their contribution to classification, quantified using the mean decrease Gini coefficient, a metric quantifying variable importance in a given model. To further distinguish if bee species shape microbial communities even when foraging on the same pollen species, we selected provisions from each species comprised by at least 50% manzanita (N=19 *O. ribifloris*, N=4 *O. lignaria* samples) and compared bacterial and fungal diversity and composition between this subset using linear models and PerMANOVA as above.

To examine the role of geographic distance in explaining variation in provision composition, we tested for congruence in distance decay relationships using Mantel correlograms for pollen, bacteria and fungi. We used Mantel tests implemented in the vegan package to determine if similarity in pollen composition was associated with variation in bacterial or fungal composition. To directly compare the role of geographic distance and pollen composition in predicting microbial composition, we used partial Mantel tests to assess the effect of geographic distance between sites while controlling for pollen composition or vice versa (Goslee & Urban 2007). We repeated partial Mantel tests for each bee species separately to examine if patterns were consistent within bee species.

Finally, we examined if the occurrence and abundance of pathogenic or symbiotic genera including *Ascosphaera*, *Sodalis,* and *Wolbachia* varied between bee species or across sites. We used binomial and linear models to predict if the presence or relative abundance of all ASVs assigned to focal genera differed by bee species, site, or their interaction. To examine geographic structuring for each, we estimated Moran’s I (Bjornstad 2016) for the relative sequence abundance of each focal genera using data from the bee species in which the microbe group was detected. We also assessed if pollen composition within a nest was associated with the prevalence of each focal microbial group using PerMANOVA implemented in vegan (Oksanen *et al*. 2007).

## Results

### Bee species and sites host divergent pollen composition

Pollen diversity and composition in bee provisions varied by bee species and geographic site (Table 1, Fig 2). The degree of turnover in each of these communities among sites also differed between bee species (significant site x species interaction; Table 1, Fig 2). Provisions of *O. lignaria* showed much greater turnover among sites than did those of *O. ribifloris* (Fig 3). Pollen generic richness varied among sites, which ranged from 1-16 genera detected in a provision. Unsurprisingly, the provisions made by the diet specialist *O. ribifloris* had lower pollen richness compared to the generalist *O. lignaria* (F_1,105_=53, p=0.01). Also as expected, *O. ribifloris* provisions were overwhelmingly comprised of pollens from the Ericaceae including the genus *Arctostaphylos* in unmanaged sites and *Arbutus* in agricultural sites (Fig 3). In contrast, *O. lignaria* pollen composition varied strongly among sites. Certain sites were dominated by the genera *Cercis*, *Acer* or *Hydrophyllum*, whereas others exhibited a mixed composition including use of *Arctostaphylos* and *Arbutus*.

**Figure 3.**
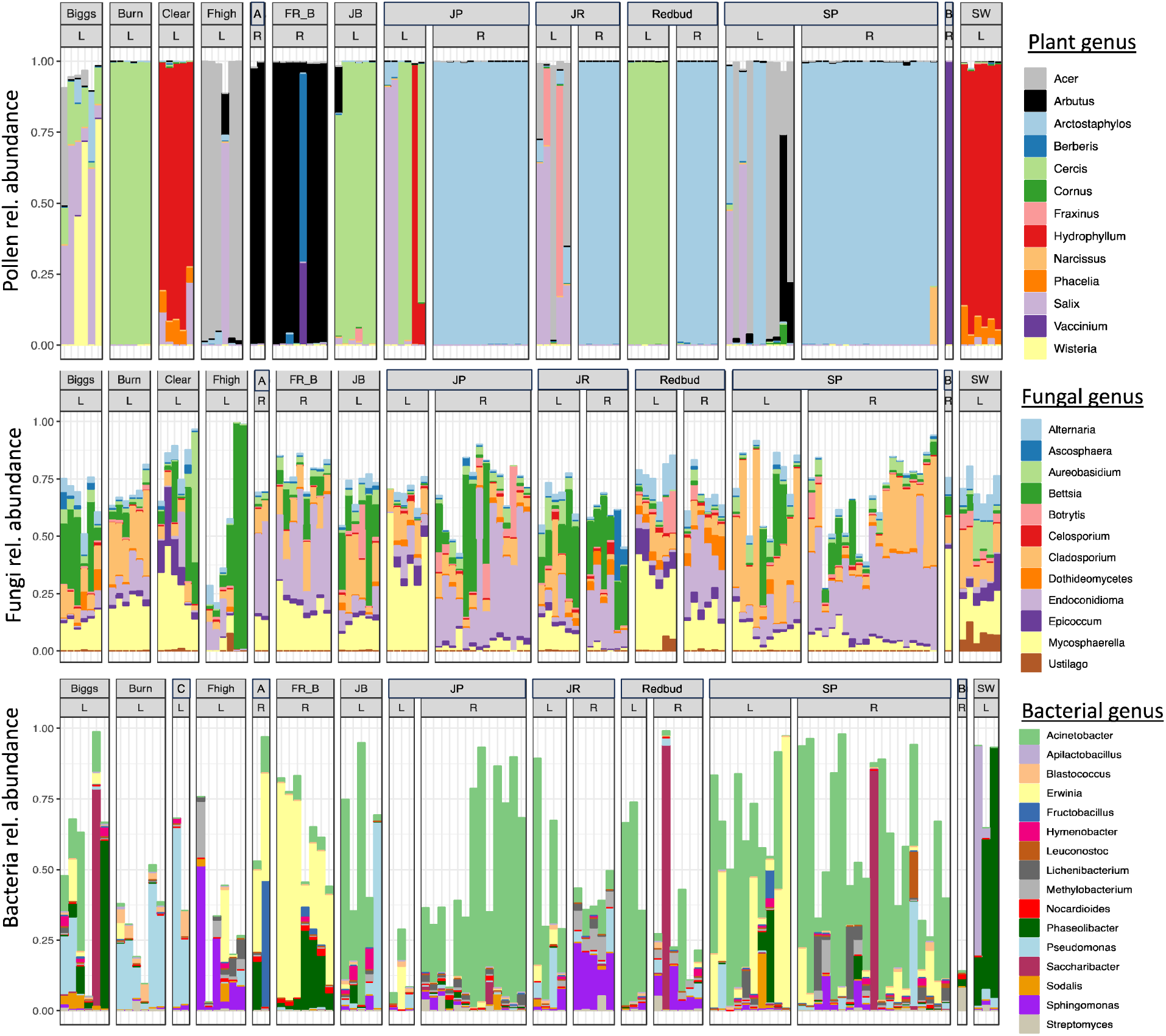
Composition of A) Pollen (rbcL), B) Fungi (ITS), and C) Bacteria (16S) detected in *Osmia ribifloris* (R) *and O. lignaria* (L) provisions using amplicon sequencing. Relative abundance is visualized as the proportion of reads in a sample that were annotated to the focal genus (top 20 visualized, in some cases multiple ASVs within a given genus were most abundant and are collapsed for readability). The top heading in each panel indicates sampling site and within each, individual bars indicate single provisions from each bee species sampled at a given site. Because pollen composition is often not proportional to the input pollen composition due to differential extraction or amplification, we used presence/absence of a genus in all community (Jaccard) and diversity analyses. Sample sizes are in some cases unequal across amplicon datasets due to sequencing failure for a given gene region. Abbreviations for some sites were shortened, here C indicates “Clear”; A for “FR_A” and B for Stockton-Bix.

**Table 1.** PerMANOVA results examining the explanatory power of bee species identity, geographic sampling site and their interaction in explaining variation in pollen species composition, fungal community composition and bacterial community composition.

| Predictor | R<sup>2</sup> | F-value | P-value |
| --- | --- | --- | --- |
| **Bee species** | 0.083 | 16.22 | 0.001 |
| **Geographic site** | 0.33 | 5.34 | 0.001 |
| **Species x Site** | 0.058 | 3.76 | 0.001 |
| **<u>Fungi</u>** |  |  |  |
| **Bee species** | 0.12 | 25.58 | 0.001 |
| **Geographic site** | 0.3 | 5.18 | 0.001 |
| **Species x Site** | 0.059 | 3.99 | 0.001 |
| **<u>Bacteria</u>** |  |  |  |
| **Bee species** | 0.026 | 3.59 | 0.001 |
| **Geographic site** | 0.29 | 3.23 | 0.001 |
| **Species x Site** | 0.05 | 2.59 | 0.001 |

### Bee species and sites differ in microbial composition

Similarly, the fungal and bacterial composition of provisions varied between bee species and among sites (Table 1, Fig 2). The degree of turnover among sites also differed between bee species with *O. lignaria* showing less differentiation among sites relative to *O. ribifloris* (significant site x species interaction; Table 1) (Fig 2, Figs S2-S3). The alpha diversity of microbes in provisions did not differ significantly between bee species (fungi F_1,106_ =3.38, p=0.07; bacteria F_1,87_=1.91, p=0.16). Fungal ASV richness varied among sites (F_12,106_=3.02, p=0.001), but bacterial ASV richness did not differ among sites (F_12,87_=1.31, p=0.22). Across bee species, pollen generic richness positively associated with increasing fungal richness (t=2.62, p=0.01), but not with bacterial richness (t=-0.14, p=0.88).

We examined if any microbial taxa formed a core microbiome across provisions of both bee species and found no microbial genus common to all provisions. The fungal genera *Bettsia* and *Mycosphaerella* occurred in 80% (but not 100%) of provisions with at least 1% relative abundance, but no bacterial genera occurred in more than 50% of provisions at 1% relative abundance. Nonetheless, specific microbial genera characterized each bee species’ stored food, meaning that although not all provisions of a given species contained these genera, each were detected at higher abundance in provisions of one bee species compared to the other. Provisions of the oligolectic bee *O. ribifloris* hosted higher relative abundance of the typically floral-associated genera *Acinetobacter* and *Zymobacter*, *Saccharibacter* and the Actinomycetes *Knoellia* and *Streptomyces* (Fig S2; DESeq2 all false discovery rate (FDR)<0.05). The polylectic bee *O. lignaria* hosted a greater relative abundance of the insect symbiont *Wolbachia*, as well as *Rubrobacter*, *Pantoea*, *Sodalis* and other bacteria and fungi across many orders including the putative plant pathogen *Alternaria* (Fig S3). To assess the extent to which microbial composition in the provision could predict bee species that formed the provision, we used random forest analysis. Based on random forest analyses, the fungal microbiome correctly distinguished bee species with 93% accuracy while bacterial composition predicted bee species with 86% accuracy. The same microbial genera that exhibited significant differential abundance between species (Figs S2-S3) were often among the top features distinguishing bee species (Figs S4-S5).

To more closely examine if differential pollen use by bee species or other species-specific modifications to provisions shapes differences in microbial composition, we subset our data to provisions containing mostly manzanita from the same site (N=19 *O. ribifloris*, N=4 *O. lignaria*). In this comparison, bacterial diversity and composition were indistinguishable between bee species (bacteria diversity t-test p=0.5; composition PerMANOVA p=0.58). Similarly, neither fungal diversity nor composition differed between the two (diversity t-test p=0.70; composition PerMANOVA p=0.14). Together these outcomes suggest a primary role of pollen composition rather than species-specific provision processing in shaping microbiome composition.

### Pollen composition, not geographic proximity, predicts microbial composition in provisions

Across the full dataset, pollen composition, but not geographic distance, was significantly associated with variation in fungal and bacterial composition (Table 2). This was not due to underlying spatial structuring in pollen use by bees: pollen composition of bee provisions exhibited significant positive autocorrelation at the smallest distance class considered (within-site), but negative at intermediate distance classes, resulting in weak overall geographic structuring among populations (Table 2). Moreover, neither fungal nor bacterial composition in provisions displayed significant overall spatial autocorrelation (fungi Mantel r=0.03, p=0.16; bacteria Mantel r=0.03; p=0.25).

**Table 2.** Mantel tests examining the relationship between the composition of pollen, fungi or bacterial communities in individual bee provisions and geographic distance between sites or pollen dissimilarity among individual provisions (Jaccard). Significance is determined by comparison with N=999 randomizations, with significant associations in bold text. Partial Mantel tests compare two focal matrices conditioned on a third matrix.

| Mantel tests |  |  |  |  |
| --- | --- | --- | --- | --- |
|  | <u>Geography as a sole predictor</u> |  | <u>Pollen as a sole predictor</u> |  |
| Composition | r | p-value | r | p-value |
| Pollen | 0.027 | 0.06 | - | - |
| Fungi | 0.03 | 0.16 | **0.3** | **0.001** |
| Bacteria | 0.03 | 0.25 | **0.15** | **0.001** |
| Partial Mantel tests (all data) |  |  |  |  |
|  | <u>Geography</u><br><u>(conditioned on pollen)</u> |  | <u>Pollen</u><br><u>(conditioned on Geography)</u> |  |
|  | r | p-value | r | p-value |
| Fungi | 0.02 | 0.24 | **0.3** | **0.001** |
| Bacteria | 0.03 | 0.19 | **0.16** | **0.001** |
| Partial Mantel tests (single species) |  |  |  |  |
|  | <u>Geography</u><br><u>(conditioned on pollen)</u> |  | <u>Pollen</u><br><u>(conditioned on Geography)</u> |  |
|  | r | p-value | r | p-value |
| Fungi (*O. lignaria*) | 0.008 | 0.4 | **0.13** | **0.006** |
| Bacteria (*O. lignaria*) | **0.11** | **0.02** | **0.14** | **0.002** |
| Fungi (*O. ribifloris*) | **0.11** | **0.05** | **0.09** | **0.03** |
| Bacteria (*O. ribifloris*) | 0.06 | 0.11 | **0.34** | **0.001** |

When we examined each bee species individually, pollen composition remained a significant predictor of fungal and bacterial composition for all comparisons (Table 2), but spatial proximity explained some additional variation in bacterial composition for *O. lignaria* and fungal composition for *O. ribifloris*.

### Pathogen and putative symbiont presence vary with bee species and among sites

*Sodalis*, a bacterial genus often found as an insect symbiont, was detected in both bee species and found at all but two sites (site LRT χ^2^=19.39, p=0.02). Its occurrence among sites differed depending on the bee species considered (species x site LRT χ^2^=10.2, p=0.016) but relative abundance did not differ significantly with species or sites (F_9, 84_, p=0.28), and no geographic structuring was detected (Moran’s I, p>0.17 for all distance classes). The symbiont *Wolbachia* was only detected in *O. lignaria* and varied in occurrence (site LRT χ^2^=20.7, p=0.05) and relative abundance among sites (site F_12,87_=2.2, p=0.01), with significant positive spatial autocorrelation detected only at the smallest distance class (Moran’s I=0.13 p=0.02). The relative abundance of both *Sodalis* and *Wolbachia* was associated with variation in the pollen composition among provisions (PerMANOVA p<0.01 for both), even when site and/or bee species were included in the model.

We detected multiple species within the genus *Ascosphaera*, a group of bee-specialized fungi comprised of commensal and pathogenic species. The most detected species was *Ascosphaera apis*, followed by *A. osmophila*, both of which were present in both bee species. *Ascosphaera fusiformis, A. solina, A. subglobosa, A. atra, A. osmophila, A. torchoioi,* and *A. variegata* were also detected, mostly in *O. ribifloris* (Fig 4, Table S2). The presence of any *Ascosphaera* at >1% relative sequence abundance varied across sites (glm site p=0.03). How relative abundance varied among sites also differed between bee species (glm site x bee species interaction, p=0.02). Notably, we found that *Ascosphaera* presence among sites differed between bee species. For example, at the site where *Ascosphaera* occurrence was highest for *O. ribifloris*, it was not detected in any *O. lignaria* cells, suggesting distinct routes of acquisition for each bee species. In addition, no significant spatial structuring was detected in the relative abundance of *Ascosphaera* across sites (Moran’s I, p>0.15 for all distance classes) nor was *Ascosphaera* significantly associated with pollen composition (PerMANOVA p=0.4).

**Fig 4.**
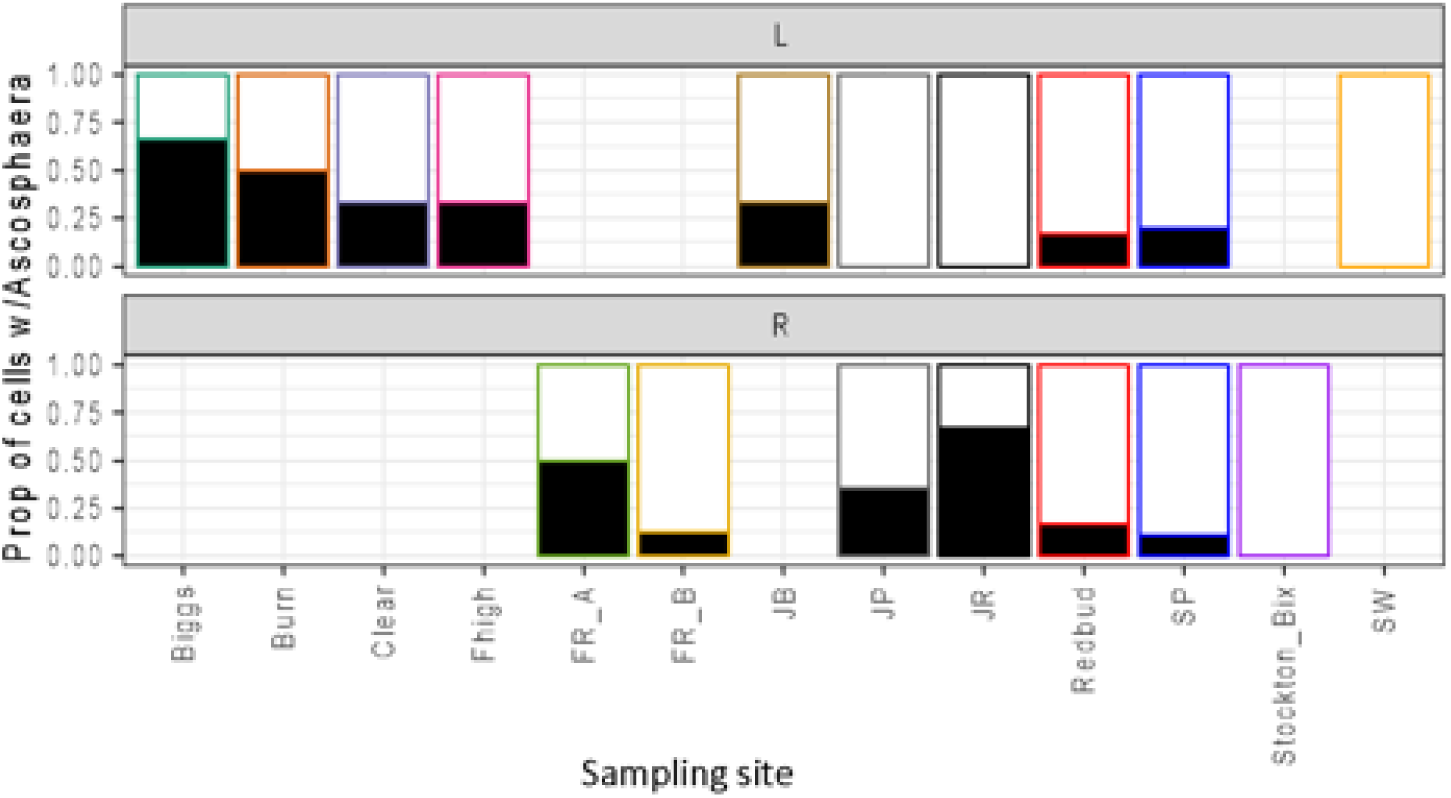
Proportion of bee provisions with *Ascosphaera* detection at 1% relative abundance or greater (depicted in black) in *O. lignaria* (L) and *O. ribifloris* (R) across geographic sampling locations. The absence of a bar at a given site indicates that the bee species was not available to be sampled at that site. The number of brood cells varied among sites, see Table S1 (FR sites had lower sample numbers). Bar outline colors match those in the map depicted in Figure 2A above.

## Discussion

Here, as in previous studies, *Osmia* populations varied in the composition of microorganisms in stored bee provisions (Rothman *et al*. 2020; Voulgari-Kokota *et al*. 2019; Westreich *et al*. 2023). By comparing pollen composition and geographic proximity across bee species and sites, we found that pollen composition explains ∼15 and 30% of variation in fungal and bacterial composition respectively, compared to only 3% explained by spatial proximity. Weak spatial effects suggest that geographic proximity, although important in explaining variation in environmental fungi and bacteria across a range of scales (Suzzi *et al*. 2023; Talbot *et al*. 2014), does not necessarily translate to the microbiome composition of pollen-sourced microbes within bees’ provisions. We suggest that floral traits may filter the bacterial and fungal communities available to bees (Herrera *et al*. 2010), as has been documented in phyllosphere and flower microbial communities (Cecala *et al*. 2024; Wang *et al*. 2023).

The finding that pollen composition predicts microbial composition in *Osmia* provisions is congruent with that from the stem-nesting bee *Ceratina* and 7 species of Megachilid bees (Voulgari-Kokota *et al*. 2019, Dew *et al*. 2020). We, like previous studies, also documented stronger links between pollen and fungi than pollen and bacteria (McFrederick & Rehan 2019b). Fungal taxa detected within provisions included *Alternaria* and *Cladosporium*, common endophytes of many flowering forb species which are transmitted via pollen (Hodgson *et al*. 2014), suggesting a mechanistic link for strong ties between fungi and pollen, although bacteria can also be pollen endophytes (Shrestha *et al*. 2024). However, significant variation in microbial composition remained unexplained; some of which may be due to variation in nectar foraging rather than to pollen. Provisions bees collect for their offspring contain 50% or more sugar by mass (Cane 2011) sourced from floral nectar, yet plant DNA from nectar is unlikely to be detected using our pollen metabarcoding approach. Floral nectar often contains abundant populations of bacteria including those common in *Osmia* provisions including *Acinetobacter*, *Apilactobacillus*, *Pseudomonas*, and *Saccharibacter* (Alvarez-Perez *et al*. 2011; Fridman *et al*. 2012; Vannette *et al*. 2021; Cecala *et al*. 2024). Given the strong overlap of such nectar associated taxa and those found in the *Osmia* provisions we sampled, we suggest that the initial species pool of microbes in *Osmia* provisions is sourced from pollen and likely the nectar of foraged flowers.

We also detected insect symbionts in provisions, including the bacteria *Wolbachia* and *Sodalis* and fungi *Ascosphaera.* For these bacterial symbionts, the primary transmission mechanism among generations is maternal inheritance (Medina Munoz *et al*. 2020; O’Neill *et al*. 1997) although environmental transmission via shared host plants may be possible (Sanaei *et al*. 2021; Tláskal *et al*. 2021). *Sodalis* and *Wolbachia* association with pollen composition may be due to environmental acquisition via ‘hub’ flowers visited by many insects (Cohen et al 2020) or differential survival in nests based on pollen composition. In contrast, the bee-specialized fungal genus *Ascosphaera* was variable among sites but not associated with pollen composition. *Ascosphaera* is thought to be transmitted to newly emerged bees via contaminated nesting materials or contact with diseased conspecifics (Stephen *et al*. 1981). Nest sanitation practices to remove infected cells may reduce the spread of this pathogen in monitored populations. In-vitro work suggests that provision-borne bacteria and fungi could suppress the growth of *Ascosphaera* (e.g. Reynaldi *et al*. 2004; Iorizzo *et al*. 2020; Christensen *et al*. 2024). Future experimental work could reveal if the presence of specific microbes, bee genetic background (Koch *et al*. 2023), environmental conditions, and other factors could contribute to population-level variation in *Ascosphaera* abundance (Gerdts *et al*. 2021).

Here we show that pollen composition is a primary determinant of the variable provision microbiome in *Osmia*, like previously documented for *Megachile* and *Ceratina* (Cohen *et al*. 2020; Nguyen & Rehan 2022) yet other bee species host consistent microbial symbionts (Christensen *et al*. 2024; Hammer *et al*. 2023). Many bee species associate with *Apilactobacillus* or other lactic acid bacteria which can dominate pollen provisions (Hammer *et al*. 2023; Kapheim *et al*. 2021; McFrederick, *et al*. 2012) and may contribute to provision preservation and larval bee survival (Nguyen & Rehan 2025). Yet the effect of *Apilactobacillus* on bees, as well as its transmission route is still unclear (Brar, *et al*. 2024). We detected *Apilactobacillus micheneri* only in a single population of *O. lignaria* in the current study, suggesting that variable exposure of *Osmia* populations to this bacterium may limit its presence rather than the inability of *Apilactobacillus* to establish in *Osmia* provisions. Investigating the host range of *Apilactobacillus*, including in species where it occurs infrequently, may further inform how bee species traits and environmental factors shape provision microbiome composition across bees more generally.

In conclusion, our work demonstrates that pollen composition, not other geographically structured factors, determine the microbiome of stored food for stem-nesting *Osmia* bee species. In these bee species, floral microbes make up the majority of microbial diversity within provisions, although we also note the variable presence of insect symbionts including *Wolbachia*, *Sodalis* and the pathogen *Ascosphaera*. We also found little influence of bee species outside of pollen choice; however, surveying a wide taxonomic range of bee species with shared pollen use could uncover variable microbial community assembly trajectories within their stored food that may be determined by bee nesting material, glandular secretions or other traits. Our study clearly links microbiome structure in plants and bees, suggesting that changes to floral microbiome composition, due to factors such as climate change or microbial biocontrol, may also influence food stability and pathogen exposure for some solitary bees.

## Author contributions

RLV and NMW planned the study, NMW, SSP and ANM conducted field work; ANM and MH performed lab work. RLV performed bioinformatic and statistical analyses, wrote the first draft of the manuscript, and all authors contributed to revisions.

## Acknowledgements

We thank the Vannette lab including Dino Sbardellati, Shawn Christensen and Jacob Cecala for comments on earlier versions of the manuscript, Madeline Handy for lab assistance, the Dalhousie IMR facility for assistance with amplicon sequencing and the Orchard Bee Association for discussion and comments on this study. This work was supported by a National Science Foundation award DEB # 1929516 to RLV, an NSF Graduate Research Fellowship to ANM and USDA Small Business Innovation award to SP, NMW and RLV. The authors declare no conflict of interest.

## Data Availability

Data are uploaded as supplementary information and will be fully accessible via NCBI, Dryad and Zenodo upon acceptance.

## References

Anderson, K.E., Carroll, M.J., Sheehan, T., Mott, B.M., Maes, P. & Corby-Harris, V. (2014). Hive-stored pollen of honey bees: many lines of evidence are consistent with pollen preservation, not nutrient conversion. Molecular Ecology, 23, 5904–5917.

Anderson, K.E., Sheehan, T.H., Mott, B.M., Maes, P., Snyder, L., Schwan, M.R., et al. (2013). Microbial Ecology of the Hive and Pollination Landscape: Bacterial Associates from Floral Nectar, the Alimentary Tract and Stored Food of Honey Bees (Apis mellifera). PLOS ONE, 8, e83125.

Bell, K. & Brosi, B. (2016). rbcL reference library.

Bjornstad, O.N. (2016). Package ‘ncf.’ Spatial nonparametric covariance functions, 5, 1.1-7.

Brar, Gagandeep, Floden, Madison, McFrederick, Quinn, Rajamohan, Arun, Yocum, George, & Bowsher, Julia. (2024). Environmentally acquired gut-associated bacteria are not critical for growth and survival in a solitary bee, Megachile rotundata. Applied and Environmental Microbiology, 90, e02076–23.

Callahan, B.J., McMurdie, P.J., Rosen, M.J., Han, A.W., Johnson, A.J.A. & Holmes, S.P. (2016). DADA2: High-resolution sample inference from Illumina amplicon data. Nat Methods, 13, 581–583.

Cambronero-Heinrichs JC, Matarrita-Carranza B, Murillo-Cruz C, Araya-Valverde E, Chavarría M, Pinto-Tomás AA. Phylogenetic analyses of antibiotic-producing *Streptomyces* sp. isolates obtained from the stingless-bee *Tetragonisca angustula* (Apidae: Meliponini). Microbiology. 2019 Mar;165(3):292–301. doi: 10.1099/mic.0.000754

Cane, J.H., Gerdin, S. & Wife, G. (1983). Mandibular Gland Secretions of Solitary Bees (Hymenoptera: Apoidea): Potential for Nest Cell Disinfection. Journal of the Kansas Entomological Society, 56, 199–204.

Cecala, J., Landucci, L. & Vannette, R. (2024). Seasonal assembly of nectar microbial communities across angiosperm plant species: assessing contributions of climate and plant traits.

Christensen, S.M., Srinivas, S.N., McFrederick, Q.S., Danforth, B.N., Buchmann, S.L. & Vannette, R.L. (2024). Symbiotic bacteria and fungi proliferate in diapause and may enhance overwintering survival in a solitary bee. The ISME Journal, 18, wrae089.

Cohen, H., McFrederick, Q.S. & Philpott, S.M. (2020). Environment Shapes the Microbiome of the Blue Orchard Bee, Osmia lignaria. Microb Ecol, 80, 897–907.

Danforth, B.N., Minckley, R.L. & Neff, J.L. (2019). The Solitary Bees: Biology, Evolution, Conservation. Princeton University Press.

Davis, N.M., Proctor, D.M., Holmes, S.P., Relman, D.A. & Callahan, B.J. (2018). Simple statistical identification and removal of contaminant sequences in marker-gene and metagenomics data. Microbiome, 6, 226.

Dew, R.M., McFrederick, Q.S. & Rehan, S.M. (2020). Diverse Diets with Consistent Core Microbiome in Wild Bee Pollen Provisions. Insects, 11, 499.

Dharampal, P.S., Carlson, C., Currie, C.R. & Steffan, S.A. (2019). Pollen-borne microbes shape bee fitness. Proc Biol Sci, 286, 20182894.

Evison, S.E. & Jensen, A.B. (2018). The biology and prevalence of fungal diseases in managed and wild bees. *Current Opinion in Insect Science*, Ecology • Parasites/Parasitoids/Biological control, 26, 105–113.

Fridman, S., Izhaki, I., Gerchman, Y. and Halpern, M., 2012. Bacterial communities in floral nectar. Environmental Microbiology Reports, 4(1), pp.97–104.

Gerdts, J.R., Roberts, J.M.K., Simone-Finstrom, M., Ogbourne, S.M. & Tucci, J. (2021). Genetic variation of *Ascosphaera apis* and colony attributes do not explain chalkbrood disease outbreaks in Australian honey bees. Journal of Invertebrate Pathology, 180, 107540.

Gilliam, M., Prest, D.B. & Lorenz, B.J. (1989). Microbiology of pollen and bee bread : taxonomy and enzymology of molds. Apidologie, 20, 53–68.

Gilliam, M., Taber, S., Lorenz, B.J. & Prest, D.B. (1988). Factors affecting development of chalkbrood disease in colonies of honey bees, *Apis mellifera*, fed pollen contaminated with *Ascosphaera apis*. Journal of Invertebrate Pathology, 52, 314–325.

Goslee, S. C., & Urban, D. L. (2007). The ecodist Package for Dissimilarity-based Analysis of Ecological Data. Journal of Statistical Software, 22(7), 1–19. 10.18637/jss.v022.i07

Hammer, T.J., Kueneman, J., Argueta-Guzmán, M., McFrederick, Q.S., Grant, Lady, Wcislo, W., et al. (2023). Bee breweries: The unusually fermentative, lactobacilli-dominated brood cell microbiomes of cellophane bees. Front. Microbiol., 14.

Herrera, C.M., Canto, A., Pozo, M.I. & Bazaga, P. (2010). Inhospitable sweetness: nectar filtering of pollinator-borne inocula leads to impoverished, phylogenetically clustered yeast communities. Proc Biol Sci, 277, 747–754.

Herrera, J., Kramer, C.L. & Reichman, O.J. (1997). Patterns of fungal communities that inhabit rodent food stores: effect of substrate and infection time. Mycologia, 89, 846–857.

Hodgson, S., de Cates, C., Hodgson, J., Morley, N.J., Sutton, B.C. & Gange, A.C. (2014). Vertical transmission of fungal endophytes is widespread in forbs. Ecology and Evolution, 4, 1199–1208.

Iorizzo, M., Lombardi, S.J., Ganassi, S., Testa, B., Ianiro, M., Letizia, F., Succi, M., Tremonte, P., Vergalito, F., Cozzolino, A. and Sorrentino, E., 2020. Antagonistic activity against *Ascosphaera apis* and functional properties of *Lactobacillus kunkeei* strains. Antibiotics, 9(5), p.262.

Kapheim, K.M., Johnson, M.M. & Jolley, M. (2021). Composition and acquisition of the microbiome in solitary, ground-nesting alkali bees. Sci Rep, 11, 2993.

Koch, J.B.U., Branstetter, M.G., Cox-Foster, D.L., Knoblett, J., Lindsay, T.-T.T., Pitts-Singer, T.L., et al. (2023). Novel Microsatellite Markers for Osmia lignaria (Hymenoptera: Megachilidae): A North American Pollinator of Agricultural Crops and Wildland Plants. Journal of Insect Science, 23, 1.

Li, Y., Liu, S., Zhang, S., Liu, T., Qin, H., Shen, C., Liu, H., Yang, F., Yang, C., Yin, Q. and Mao, J., 2022. Spatiotemporal distribution of environmental microbiota in spontaneous fermentation workshop: The case of Chinese Baijiu. Food Research International, 156, p.111126.

Liaw, A. & Wiener, M. (2002). Classification and regression by randomForest. R news, 2, 18– 22.

Louw, N.L., Lele, K., Ye, R., Edwards, C.B. & Wolfe, B.E. (2023). Microbiome Assembly in Fermented Foods. Annual Review of Microbiology, 77, 381–402.

Love, M.I., Huber, W. and Anders, S., 2014. Moderated estimation of fold change and dispersion for RNA-seq data with DESeq2. Genome biology, 15, pp.1–21.

McFrederick, Q.S. & Rehan, S.M. (2019). Wild Bee Pollen Usage and Microbial Communities Co-vary Across Landscapes. Microb Ecol, 77, 513–522.

McFrederick, Quinn S., Q.S., Wcislo, W.T., Taylor, D.R., Ishak, H.D., Dowd, S.E. & Mueller, U.G. (2012). Environment or kin: whence do bees obtain acidophilic bacteria? Molecular Ecology, 21, 1754–1768.

Medina Munoz, M., Spencer, N., Enomoto, S., Dale, C. & Rio, R.V.M. (2020). Quorum sensing sets the stage for the establishment and vertical transmission of Sodalis praecaptivus in tsetse flies. PLoS Genet, 16, e1008992.

Miller, E.R., Kearns, P.J., Niccum, B.A., O’Mara Schwartz, J., Ornstein, A. and Wolfe, B.E., 2019. Establishment limitation constrains the abundance of lactic acid bacteria in the Napa cabbage phyllosphere. Applied and environmental microbiology, 85(13), pp.e00269–19.

Minckley, R.L. & Danforth, B.N. (2019). Sources and frequency of brood loss in solitary bees. Apidologie, 50, 515–525.

Morrison Whittle, P. and Goddard, M.R., 2018. From vineyard to winery: a source map of microbial diversity driving wine fermentation. Environmental microbiology, 20(1), pp.75–84.

Nguyen, P.N. & Rehan, S.M. (2022). Developmental microbiome of the small carpenter bee, Ceratina calcarata. Environmental DNA, 4, 808–819.

Nguyen, P.N. & Rehan, S.M. (2023). Wild bee and pollen microbiomes across an urban–rural divide. FEMS Microbiology Ecology, 99, fiad158.

Nilsson, R.H., Larsson, K.-H., Taylor, A.F.S., Bengtsson-Palme, J., Jeppesen, T.S., Schigel, D., et al. (2019). The UNITE database for molecular identification of fungi: handling dark taxa and parallel taxonomic classifications. Nucleic Acids Research, 47, D259–D264.

O’Neill, S.L., Hoffman, A.A. & Werren, J.H. (1997). Influential passengers: inherited microorganisms and arthropod reproduction.

Oksanen, J., Kindt, R., Legendre, P., O’Hara, B., Stevens, M.H.H., Oksanen, M.J. and Suggests, M.A.S.S., 2007. The vegan package. Community ecology package, 10(631-637), p.719.

Paludo, C.R., Menezes, C., Silva-Junior, E.A., Vollet-Neto, A., Andrade-Dominguez, A., Pishchany, G., et al. (2018). Stingless Bee Larvae Require Fungal Steroid to Pupate. Sci Rep, 8, 1122.

Paul Ross, R., Morgan, S. & Hill, C. (2002). Preservation and fermentation: past, present and future. *International Journal of Food Microbiology*, Notermans Special Issue, 79, 3–16.

Phuong N Nguyen, Sandra M Rehan, Supporting wild bee development with a bacterial symbiont, *Journal of Applied Microbiology*, Volume 136, Issue 1, January 2025, lxae317, 10.1093/jambio/lxae317

R Core Team. (2021). R: A language and environment for statistical computing.

Reynaldi, F.J., De Giusti, M.R. and Alippi, A.M., 2004. Inhibition of the growth of Ascosphaera apis by Bacillus and Paenibacillus strains isolated from honey. Revista Argentina de Microbiologia, 36(1), p.52.

Rothman, J.A., Cox-Foster, D.L., Andrikopoulos, C. & McFrederick, Q.S. (2020). Diet Breadth Affects Bacterial Identity but Not Diversity in the Pollen Provisions of Closely Related Polylectic and Oligolectic Bees. Insects, 11, 645.

Sampson, B.J., Stringer, S.J. & Marshall, D.A. (2013). Blueberry Floral Attributes and Their Effect on the Pollination Efficiency of an Oligolectic Bee, Osmia ribifloris Cockerell (Megachilidae: Apoidea). HortScience, 48, 136–142.

Sanaei, E., Charlat, S. & Engelstädter, J. (2021). Wolbachia host shifts: routes, mechanisms, constraints and evolutionary consequences. Biological Reviews, 96, 433–453.

Shrestha, A., Limay-Rios, V., Brettingham, D.J.L. & Raizada, M.N. (2024). Maize pollen carry bacteria that suppress a fungal pathogen that enters through the male gamete fertilization route. Front. Plant Sci., 14.

Stephen, W.P., Vandenberg, J.D. and Fichter, B.L., 1981. Etiology and epizootiology of chalkbrood in the leafcutter bee, *Megachile rotundata* (Fabricius), with notes on Ascosphaera species. Utah Agricultural Experiment Station Bulletin, 653, p.1.

Steffan, S.A., Dharampal, P.S., Danforth, B.N., Gaines-Day, H.R., Takizawa, Y. & Chikaraishi, Y. (2019). Omnivory in Bees: Elevated Trophic Positions among All Major Bee Families. Am Nat, 194, 414–421.

Steffan, S.A., Dharampal, P.S., Kueneman, J.G., Keller, A., Argueta-Guzmán, M.P., McFrederick, Q.S., et al. (2024). Microbes, the ‘silent third partners’ of bee–angiosperm mutualisms. Trends in Ecology & Evolution, 39, 65–77.

Stothart, M.R., Spina, H.A., Hotchkiss, M.Z., Ko, W. & Newman, A.E.M. (2023). Seasonal dynamics in the mammalian microbiome between disparate environments. Ecology and Evolution, 13, e10692.

Suzzi, A.L., Stat, M., Gaston, T.F. & Huggett, M.J. (2023). Spatial patterns in host-associated and free-living bacterial communities across six temperate estuaries. FEMS Microbiology Ecology, 99, fiad061.

Talbot, J.M., Bruns, T.D., Taylor, J.W., Smith, D.P., Branco, S., Glassman, S.I., et al. (2014). Endemism and functional convergence across the North American soil mycobiome. Proceedings of the National Academy of Sciences, 111, 6341–6346.

Tláskal, V., Pylro, V.S., Žifčáková, L. & Baldrian, P. (2021). Ecological Divergence Within the Enterobacterial Genus Sodalis: From Insect Symbionts to Inhabitants of Decomposing Deadwood. Front. Microbiol., 12.

Torchio, P.F. (1989). In-Nest Biologies and Development of Immature Stages of Three Osmia Species (Hymenoptera: Megachilidae). Annals of the Entomological Society of America, 82, 599–615.

Vannette, R.L., McMunn, M.S., Hall, G.W., Mueller, T.G., Munkres, I. and Perry, D., 2021. Culturable bacteria are more common than fungi in floral nectar and are more easily dispersed by thrips, a ubiquitous flower visitor. FEMS Microbiology Ecology, 97(12), p.fiab150.

Voulgari-Kokota, A., Ankenbrand, M.J., Grimmer, G., Steffan-Dewenter, I. & Keller, A. (2019). Linking pollen foraging of megachilid bees to their nest bacterial microbiota. Ecology and Evolution, 9, 10788–10800.

Voulgari-Kokota, A., Steffan-Dewenter, I. & Keller, A. (2020). Susceptibility of Red Mason Bee Larvae to Bacterial Threats Due to Microbiome Exchange with Imported Pollen Provisions. Insects, 11, 373.

Wang, Z., Jiang, Y., Zhang, M., Chu, C., Chen, Y., Fang, S., et al. (2023). Diversity and biogeography of plant phyllosphere bacteria are governed by latitude-dependent mechanisms. New Phytologist, 1534–1547.

Westreich, L.R., Westreich, S.T. & Tobin, P.C. (2023). Bacterial and Fungal Symbionts in Pollen Provisions of a Native Solitary Bee in Urban and Rural Environments. Microb Ecol, 86, 1416–1427.

Williams, N.M. & Tepedino, V.J. (2003). Consistent mixing of near and distant resources in foraging bouts by the solitary mason bee Osmia lignaria. Behavioral Ecology, 14, 141– 149.

Wynns, A.A., 2012. The Bee Specialist Fungus Family Ascosphaeraceae and Its Allies: Systematics, Ecology and Co-evolution with Solitary Bees: Phd Thesis. Department of Agriculture and Ecology, University of Copenhagen.

