## Supplementary Information for "Pollen diet, more than geographic distance, shapes provision microbiome composition in two species of cavity-nesting bees"

**Supplementary Figures and Tables** for “Pollen diet, more that geographic distance, explains variation in the provision microbiome composition in two species of cavity-nesting bees”

**Supplementary Table 1**. Sites sampled and number of provisions sequenced for each *Osmia* species at each site. Samples were either sampled using sterile technique in the field (grooved wooden boards) or transported to the lab (nesting straws). Columns indicate which years bees were sampled, and the estimated number of nests of each species at each site.

| SiteID | *O. lignaria*  *sampled* | *O. ribifloris*  *sampled* | *Nesting substrate* | *Bees added** | *2020 sampling* | *2022 sampling* | *O. ribifloris nests in sampling period* | *O. lignaria nests in sampling period* |
| --- | --- | --- | --- | --- | --- | --- | --- | --- |
| Biggs | 6 | 0 | Grooved boards |  | X |  | 0 | 110 |
| Burn | 6 | 0 | straw |  | X |  | 0 | 282 |
| Clear | 6 | 0 | straw |  | X |  | 18 | 647 |
| Fhigh | 6 | 0 | straw |  | X |  | 1 | 33 |
| FR_A | 0 | 2 | straw | Yes |  | X | 11 | 0 |
| FR_B | 0 | 8 | straw | Yes |  | X | 11 | 0 |
| JB | 6 | 0 | straw |  | X |  | 0 | 282 |
| JP | 6 | 14 | Grooved boards |  | X | X | 1033 | 47 |
| JR | 6 | 6 | Grooved boards |  | X |  | 42 | 24 |
| Redbud | 6 | 6 | Grooved boards |  | X |  | 18 | 647 |
| SP | 10 | 20 | straw |  | X | X | 309 | 332 |
| Stockton_Bix | 0 | 1 | straw | Yes |  | X | 1 | 0 |
| SW | 6 | 0 | straw |  | X |  | 0 | 495 |

***** *if blank, bee species were nesting at the site prior to sampling year*

**Supplementary Table S2.** *Ascosphaera* species detected using ITS amplicon sequencing within each bee species. Numbers in each cells indicates the proportion of sites sampled where a given actual sequence variant (ASV) was detected within that species.

| *Ascosphaera* species | *O. lignaria* | *O. ribifloris* |
| --- | --- | --- |
| *A. apis* | 0.5 | 0.83 |
| *A. fusiformis-ASV1* | 0 | 0.14 |
| *A. fusiformis-ASV2* | 0.2 | 0.00 |
| *A. solina-ASV1* | 0 | 0.14 |
| *A. solina-ASV2* | 0 | 0.29 |
| *A. subglobosa-ASV1* | 0 | 0.14 |
| *A. subglobosa-ASV2* | 0.3 | 0.43 |
| *A. atra* | 0 | 0.43 |
| *A. osmophila* | 0.3 | 0.43 |
| *A. torchioi* | 0.2 | 0.57 |
| *A. variegata* | 0 | 0.14 |
| *Ascosphaera* unknown species | 0.3 | 0.14 |

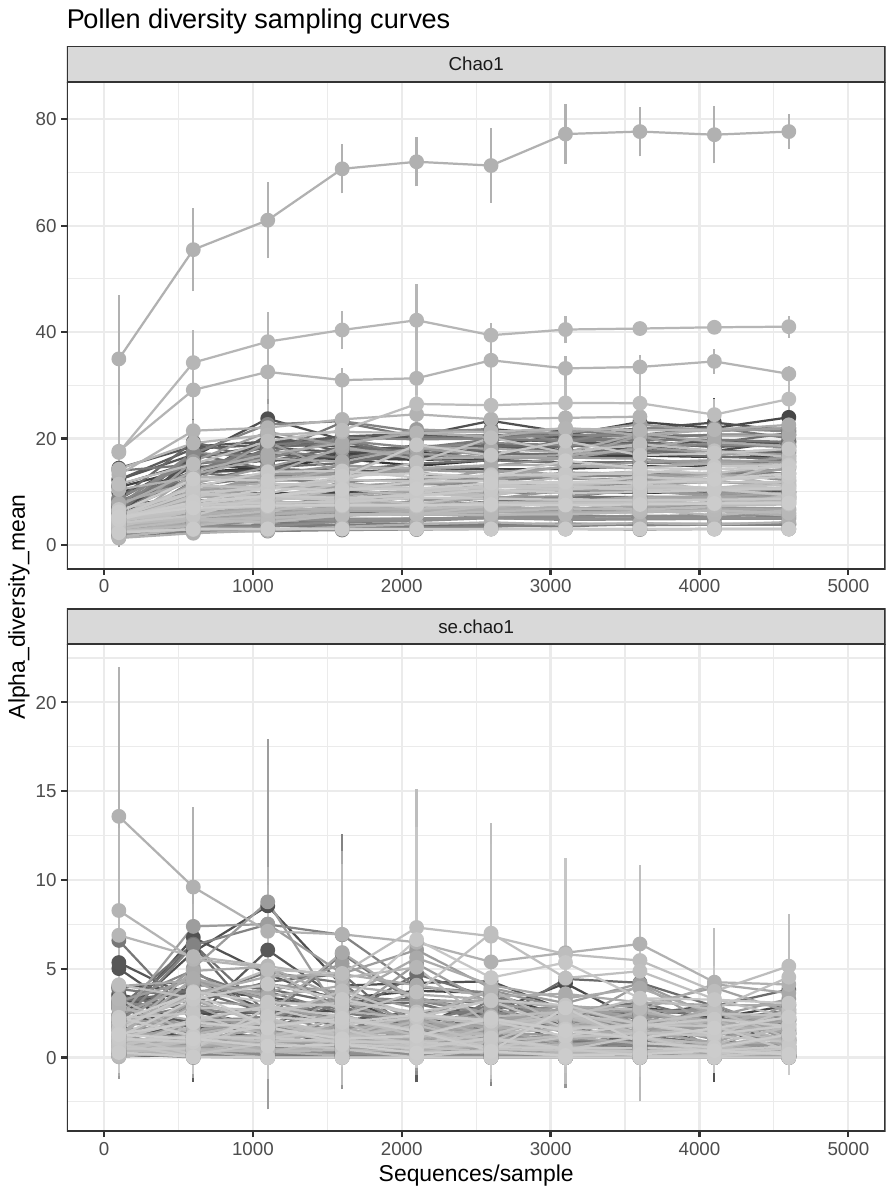

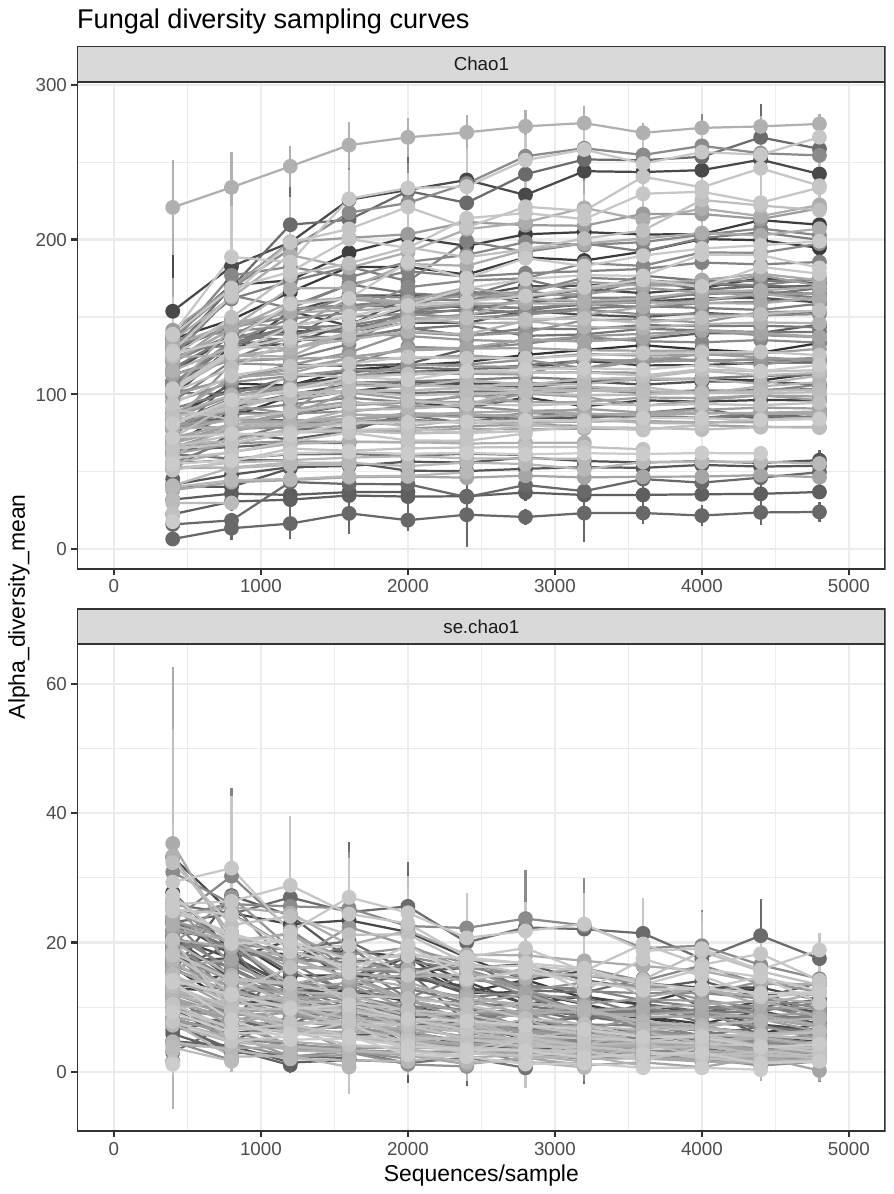

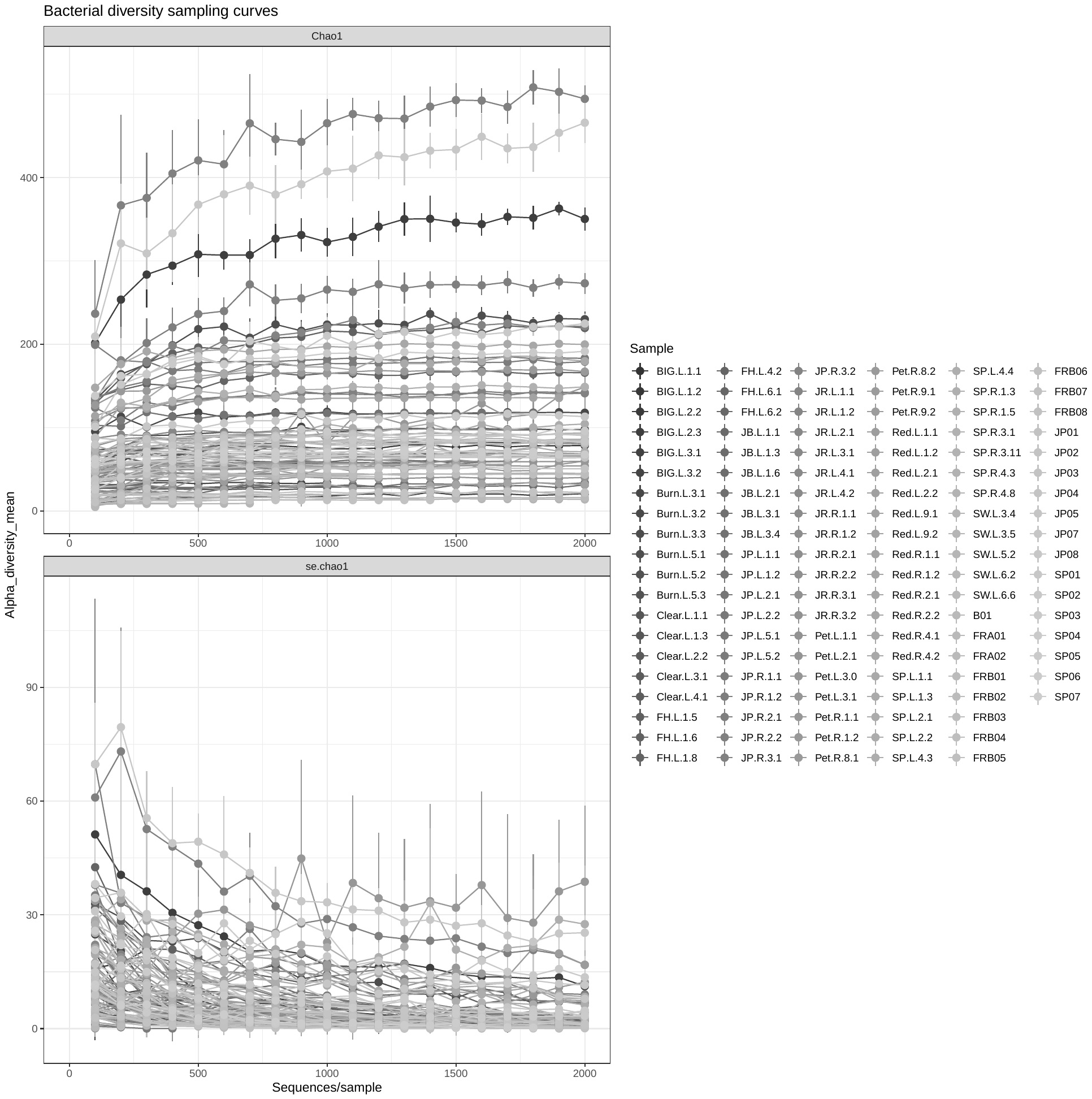

**Supplementary Figure S1** Rarefaction curves for A) Pollen, b) Fungi, and C) Bacteria within individual samples. The X-axis indicates sequencing depth and y-axis indicates # of ASVs estimated using Chao1 richness at a given sequencing depth.

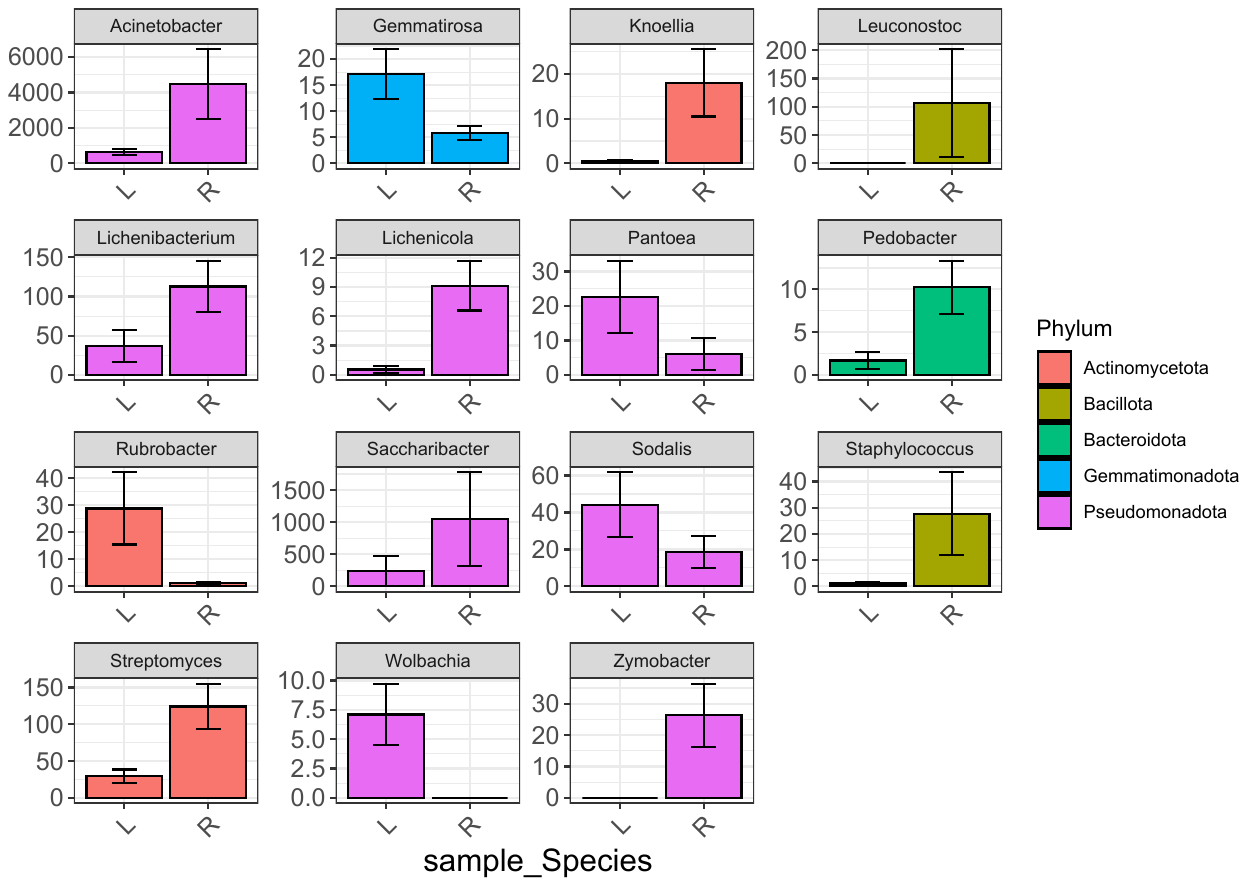

**Supplementary Figure S2.** Bacterial genera that differ in relative abundance between bee species provisions including *Osmia lignaria* (L) or *O. ribifloris* (R), as detected using DeSEQ 2 with FDR <0.05. The average abundance in each species +/-1SE is shown and bars are colored by the bacterial Phylum.

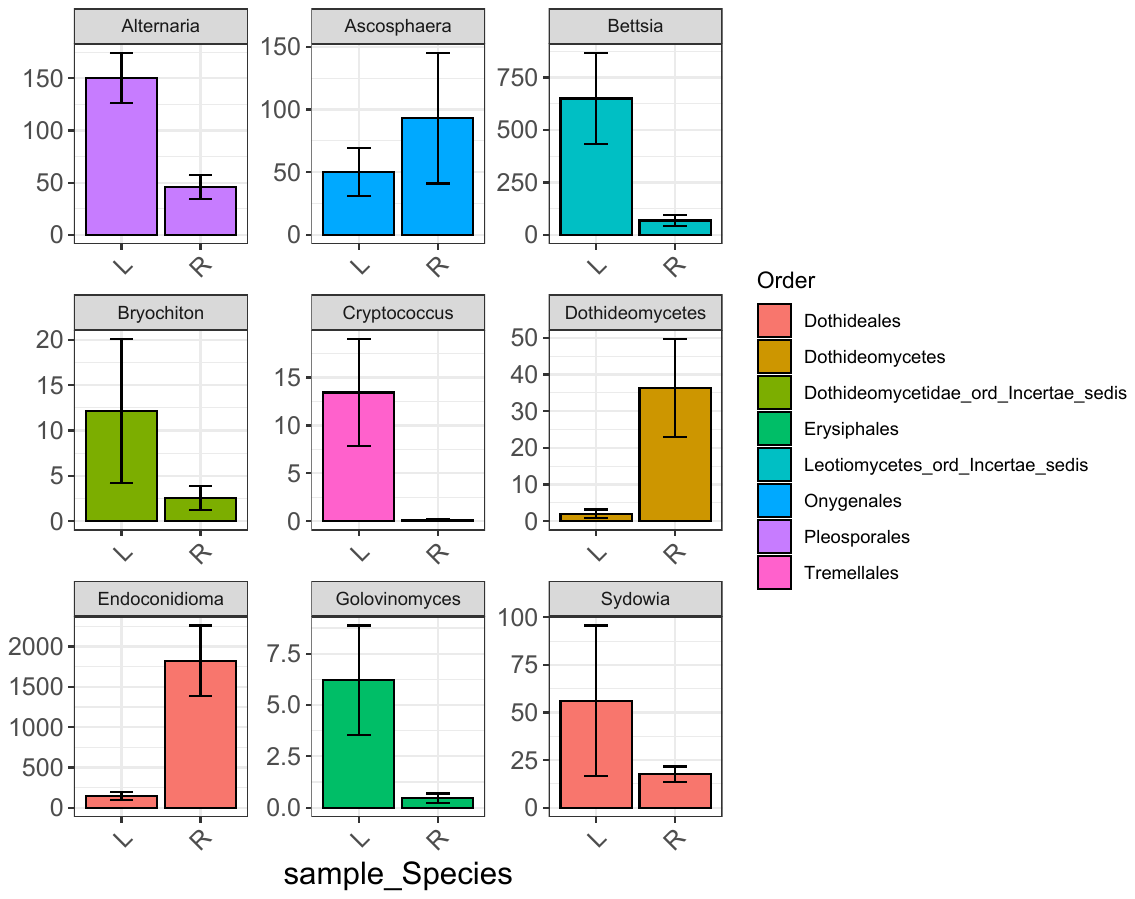

**Supplementary Figure S3.** Fungal genera that differ in relative abundance between bee species provisions including *Osmia lignaria* (L) or *O. ribifloris* (R), as detected using DESeq2 with FDR <0.05. The average abundance at each site +/-1SE is shown and bars are colored by the Order.

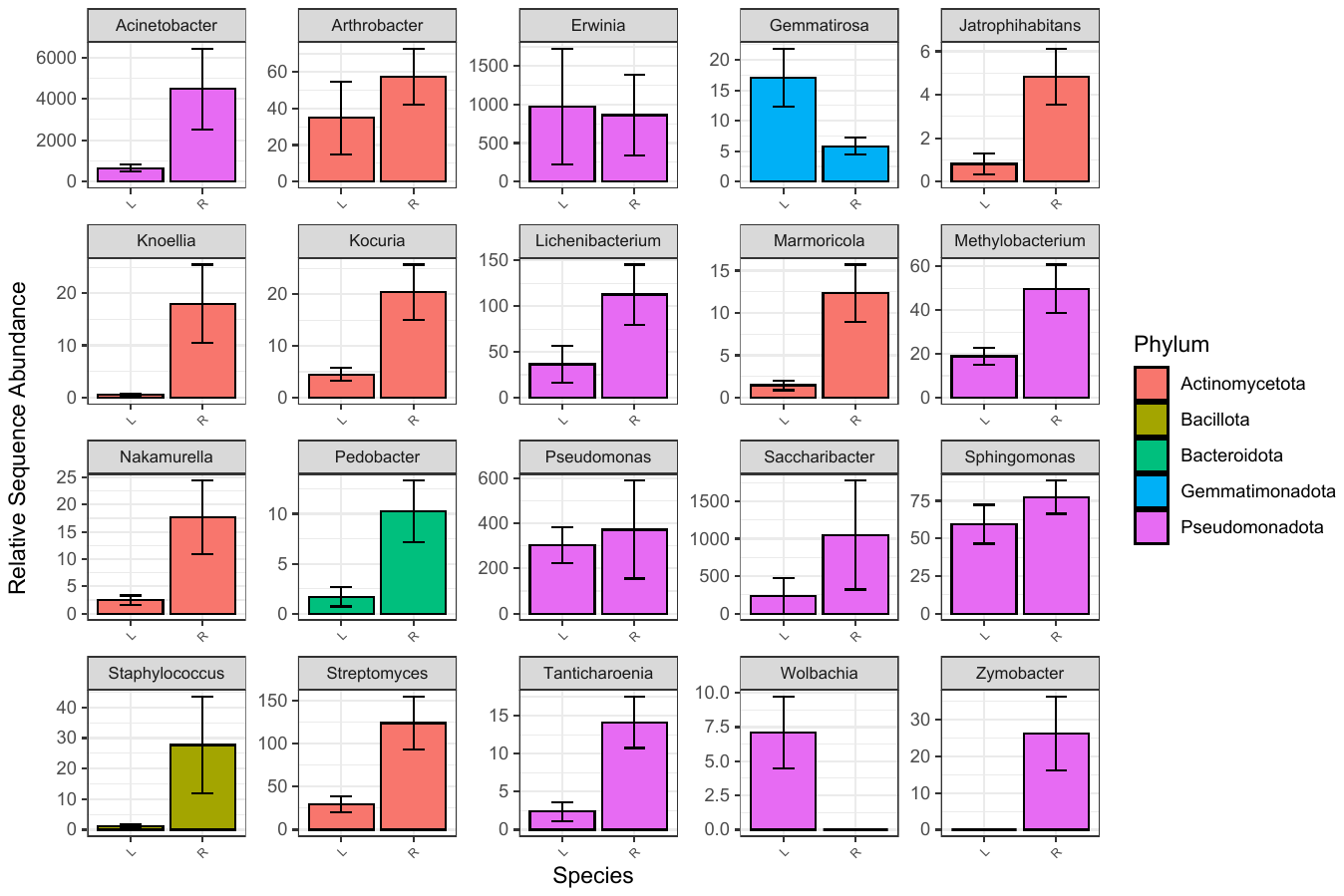

**Supplementary Figure S4.** Top 20 bacterial genera with the greatest contribution to distinguishing microbiome composition between bee species including *Osmia lignaria* (L) or *O. ribifloris* (R), measured by mean decrease in Gini in random forest models. The average abundance in each species +/- 1 SE is shown and bars are colored by the bacteria phylum.

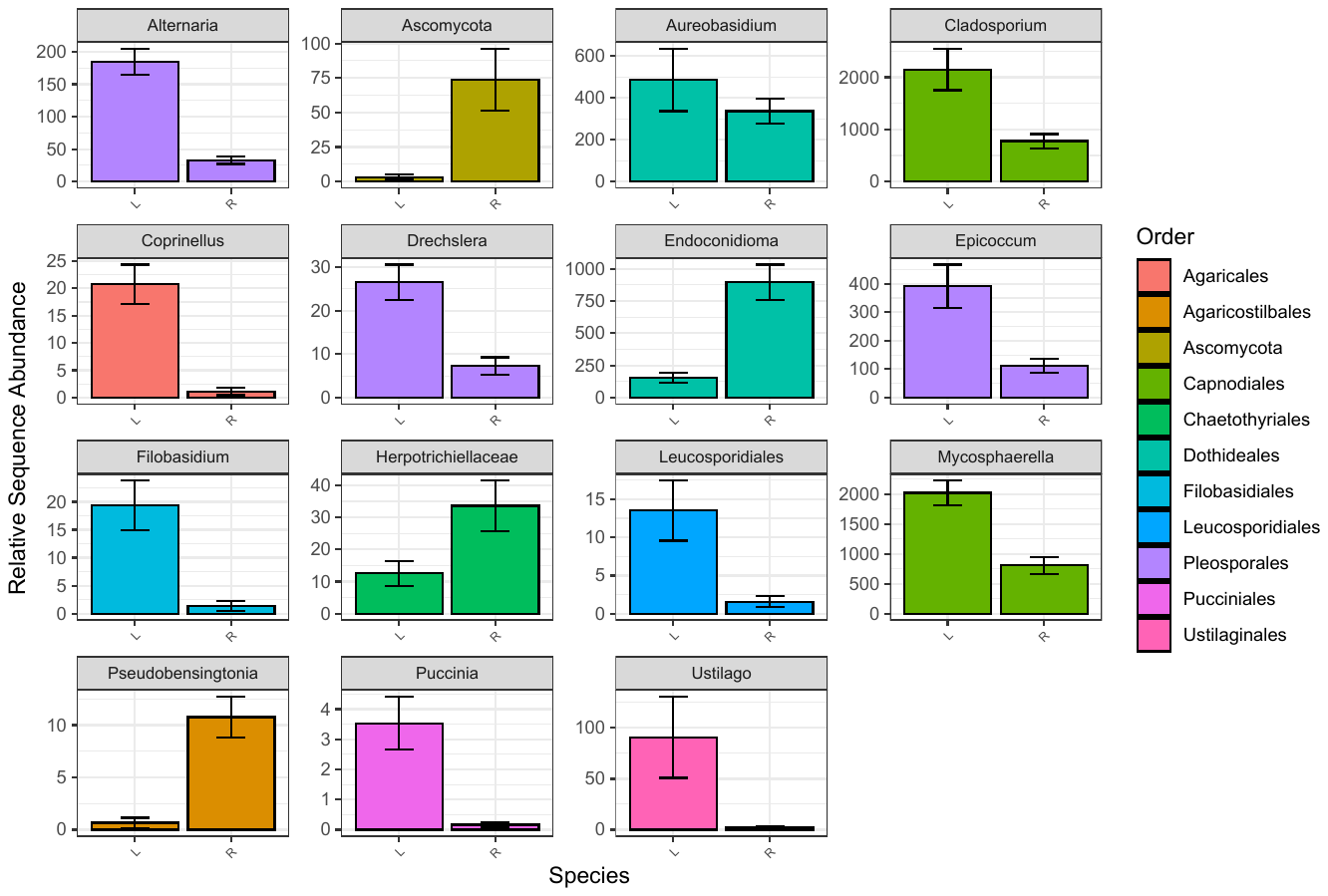

**Supplementary Figure S5.** Fungal genera with the greatest contribution to distinguishing microbiome composition between bee species including *Osmia lignaria* (L) or *O. ribifloris* (R), measured by mean decrease in Gini in random forest models. The average abundance in each species +/-1 SE is shown and bars are colored by the Order.
